# Tamoxifen Response at Single Cell Resolution in Estrogen Receptor-Positive Primary Human Breast Tumors

**DOI:** 10.1101/2023.04.01.535159

**Authors:** Hyunsoo Kim, Austin A. Whitman, Kamila Wisniewska, Rasha T. Kakati, Susana Garcia-Recio, Benjamin C. Calhoun, Hector L. Franco, Charles M. Perou, Philip M. Spanheimer

**Affiliations:** Lineberger Comprehensive Cancer Center, University of North Carolina, Chapel Hill, NC; Department of Pathology and Laboratory Medicine, University of North Carolina at Chapel Hill, Chapel Hill, NC; Department of Genetics, University of North Carolina, Chapel Hill, NC; Computational Medicine Program, University of North Carolina, Chapel Hill, NC; Department of Surgery, University of North Carolina, Chapel Hill, NC

**Author notes:** Authors contributed equally to this work (Hyunsoo Kim, Austin A. Whitman).

## Abstract

In ER+/HER2- breast cancer, multiple measures of intra-tumor heterogeneity are associated with worse response to endocrine therapy. To investigate heterogeneity in response to treatment, we developed an operating room-to-laboratory pipeline for the collection of live human tumors and normal breast specimens immediately after surgical resection for processing into single-cell workflows for experimentation and genomic analyses. We demonstrate differences in tamoxifen response by cell type and identify distinctly responsive and resistant subpopulations within the malignant cell compartment of human tumors. Tamoxifen resistance signatures from 3 distinct resistant subpopulations are prognostic in large cohorts of ER+ breast cancer patients and enriched in endocrine therapy resistant tumors. This novel ex vivo model system now provides a foundation to define responsive and resistant sub-populations within heterogeneous tumors, to develop precise single cell-based predictors of response to therapy, and to identify genes and pathways driving resistance to therapy.

## INTRODUCTION

Breast cancer is the most common cancer in women and around 600,000 women die from breast cancer worldwide each year^1^. Approximately 75% of breast cancers are categorized within the estrogen receptor positive (ER+)/luminal subtypes, which are characterized by expression of the estrogen and progesterone receptors. In general, ER+/luminal breast cancers respond to endocrine therapy targeting the estrogen receptor (ER)^2–5^. However, up to a third of early ER+ breast cancers are inherently unresponsive or develop resistance to endocrine therapy, and resistance eventually develops in all patients with metastatic disease^6–8^.

Resistance to endocrine therapy is complex and multifactorial^9^. Known mechanisms of resistance include autologous activation of estrogen receptor signaling^10–12^, alterations of transcription factors^13–16^, and upregulation of growth factor signaling pathways^17–21^. Genomic profiling of endocrine resistant tumors has identified alterations enriched in resistant tumors, including mutations in the estrogen receptor and MAPK pathways^15,22,23^, resulting in constitutive ER activity. However, these account for only a minority of cases and mechanisms have not been described for the majority of endocrine resistant tumors.

Signatures of estrogen response genes defined from the ER+ cell line MCF-7 are highly prognostic in ER+ breast cancer patients^24^, demonstrating the importance of ER transcriptional activity on tumor biology and outcomes. Similarly, in patients treated with endocrine therapy, gene expression changes, including reduced proliferation, correlate strongly with outcome^25–28^. However, bulk profiling has been used which averages the signal obtained from assorted cellular subpopulations within the tumor. These subpopulations include stromal, immune, and normal epithelial cells, which are relatively abundant in many ER+/HER2- breast cancer specimens^29^. Tumor heterogeneity within human breast cancers is associated with metastasis and worse response to treatment, including endocrine therapy^30–37^. Distinct response to treatments has been described within breast tumors in cell populations with unique gene expression profiles^38,39^. Therefore, bulk analysis that includes cell types with distinct responses could mask relevant tumor-specific response elements and low-abundance subpopulations that may contribute to resistance. In addition, non-tumor cell types with distinct responses could limit precision in defining tumor-specific transcriptional changes. Improved understanding of how distinct populations within human tumors respond to endocrine therapy as well as the heterogeneity in response within tumors is needed to reveal underlying mechanisms of resistance.

In this study, we sought to determine how subpopulations of cells within normal breast epithelium and ER+/HER2- breast tumors respond to tamoxifen treatment using an ex vivo treatment model coupled to single-cell RNA sequencing. We hypothesize that subpopulations within ER+/HER2- human breast tumors have distinct responses to tamoxifen and that discerning heterogeneity in cellular response will improve the understanding of inherent and emerging resistance to endocrine therapy.

## RESULTS

### Clinical specimens

Studying the responsiveness of ER+/luminal breast cancer cells to tamoxifen in vivo is challenging due to features of stromal cells and intra-tumoral heterogeneity. To address these challenges, we developed a novel ex vivo system for short-term culturing coupled with single-cell RNA-seq. To demonstrate the platforms, two normal breast specimens were obtained from patients undergoing reduction mammaplasty and 10 patients with clinically ER+/HER2- treatment-naïve primary breast tumors. Patient demographics and tumor characteristics are shown in Table 1. Tumor specimens were obtained from patients with invasive ductal (n=8) or lobular (n=2) histology. All tumors were strongly positive for ER and clinically negative for HER2 (IHC 0/1+ or IHC 2+ with negative FISH). IHC for Ki-67 showed positivity ranging from 7% to 35%. PAM50 subtype determined by bulk mRNA sequencing showed that 5 tumors were classified as Luminal A, 3 as Luminal B, 1 as HER2-enriched, and 1 as normal-like.

**Table 1.** Histological data for two normal specimens and 10 tumor specimens. The PAM50 molecular subtypes were determined by bulk RNA-seq profiles. (IDC: Invasive Ductal Carcinoma, ILC: Invasive Lobular Carcinoma, [*]: FISH negative, -: Not applicable).

| Specimen ID | Histology | Race | Age | ER | PR | HER2 | Grade | Stage | PAM50 | Ki67 |
| --- | --- | --- | --- | --- | --- | --- | --- | --- | --- | --- |
| Normal_01 | Normal | White | 20 | - | - | - | - | - | - | - |
| Normal_02 | Normal | Black | 22 | - | - | - | - | - | - | - |
| Tumor_01 | IDC | Black | 68 | 100% | 95% | 2+ [*] | 2 | T1c N1a | LumA | 11% |
| Tumor_02 | IDC | Other | 44 | 95% | 65% | 2+ [*] | 2 | T3 N1a | LumB | 9% |
| Tumor_03 | ILC | White | 51 | 95% | 100% | 1+ | 3 | T1c N0 | LumA | 14% |
| Tumor_04 | IDC | White | 66 | 95% | 8% | 1+ | 1 | T1C N0 | LumA | 10% |
| Tumor_05 | IDC | White | 59 | 90% | 35% | 2+ [*] | 1 | T2 N0 | LumA | 10% |
| Tumor_06 | ILC | White | 66 | 100% | 2% | 1+ | 2 | T3 N0 (i+) | LumA | 7% |
| Tumor_07 | IDC | White | 22 | 95% | 80% | 0 | 1 | T2 N0 | LumB | 11% |
| Tumor_08 | IDC | Black | 48 | 95% | 60% | 0 [*] | 3 | T2 N0 | LumB | 18% |
| Tumor_09 | IDC | White | 44 | 90% | 100% | 0 | 3 | T2 N1a | Her2E | 35% |
| Tumor_10 | IDC | White | 71 | 100% | 5% | 0 | 2 | T1c N0 | Normal-like | 12% |

### Characterization of normal breast cell populations

Two normal breast samples were obtained and processed (Fig. 1A). In total, 13,175 single cells were identified after quality control and annotated with canonical cell type markers^40^ (Fig. 1B). Clustering occurred largely by patient which may indicate biologic variability between patients, including breast density and cycling hormone status at the time of surgery, or batch effects (Fig. 1C). When comparing the treatment conditions for each specimen, there was strong cluster overlap of tamoxifen treated cells within the fibroblast, basal epithelial, endothelial, and immune cells compared to luminal progenitor cells, indicating a more robust change in gene expression in response to tamoxifen treatment in that subpopulation.

**Figure 1.**
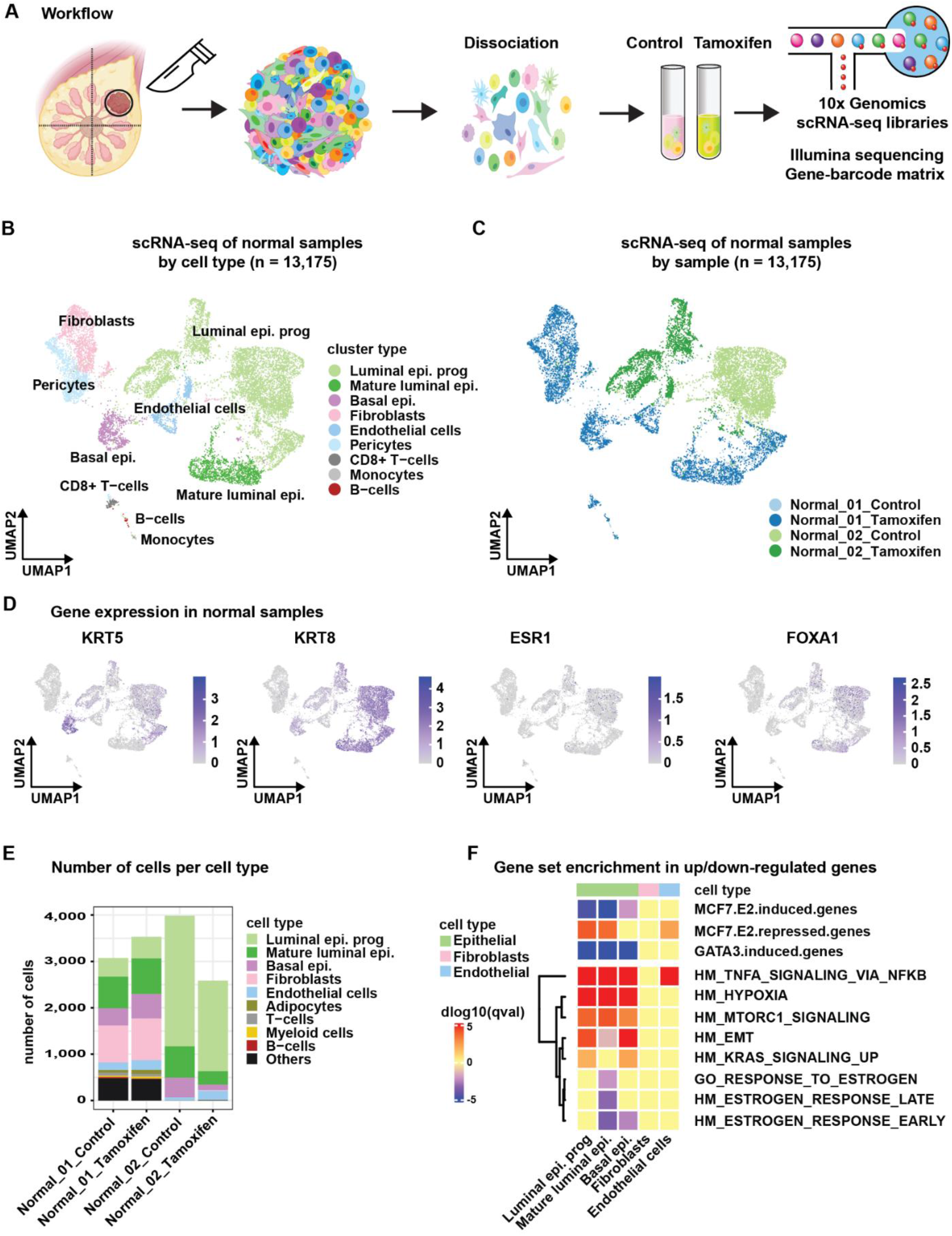
Characterization of cell type specific response to tamoxifen in normal human breast tissue. (A) Graphical representation of operating room to single-cell sequencing workflow. The breast schematic was created using BioRender.com. (B) UMAP plot of all scRNA-seq cells from two normal breast tissue samples. Cells were color-coded by cell type. (C) UMAP plot of scRNA-seq cells color-coded by sample and treatment condition. (D) Feature plots showing the expression of select epithelial markers in normal breast cells. (E) Bar chart comparing the total number of normal breast cells sequenced per treatment condition, color-coded by cell type. (F) Heatmap of gene sets enriched and depleted in differentially regulated genes in tamoxifen-treated cells relative to control cells by cell type demonstrating distinct biologic activity of tamoxifen in different cell compartments. Gene set enrichment was determined using enricher and p-values reported after correction for false discovery.

Basal cytokeratins (KRT5/14) and luminal cytokeratins (KRT8/18) were enriched in the expected cluster (Fig. 1D), demonstrating robust classification of epithelial cell types. *ESR1* transcripts, encoding the estrogen receptor, were not detected in most cells including luminal epithelial cells. *FOXA1*, a well-established pioneer transcription factor, was localized to mature and progenitor luminal epithelial cells, consistent with existing literature. Variability was seen in cell type abundance between the two specimens but did not vary significantly with tamoxifen treatment (Fig. 1E).

To evaluate the effect of time in suspension on cell subpopulation abundance, we compared an immediately created library from sample Normal_01 to the control sample after 12 hours in suspension. UMAP plots were used to visualize the data by cell type (Extended Data Fig. 1A). Cell type was the primary driver of clustering rather than time in suspension (Extended Data Fig. 1B) and canonical markers demonstrated robust cell type identification (Extended Data Fig. 1C). Fewer cells were sequenced after time in suspension but no systemic enrichment or depletion of cell populations was observed (Extended Data Fig. 1D). Differentially regulated genes showed enrichment of inflammatory signaling pathways, epithelial-mesenchymal transition (EMT) genes, and genes overlapping with estrogen response (Extended Data Fig. 1E), (complete list in Supplementary Table 1).

### Cell type specific response to tamoxifen in normal breast tissue

To determine tamoxifen-induced gene expression changes in the distinct cell subpopulations that comprise normal human breast tissue, we performed differential gene expression analysis by cell type comparing tamoxifen and control-treated cells. Upregulated and downregulated genes with tamoxifen treatment were identified separately for basal epithelial cells (BEp), luminal progenitor cells (LEp_prog), and mature luminal cells (LEp), (Supplementary Tables 2-4). Luminal progenitor cells demonstrated the most robust response by number of differentially regulated genes. To determine common and unique tamoxifen-regulated genes, we performed overlap analysis which showed the majority of genes were cell type specific. Twenty-six down-regulated genes were common to all 3 cell types (Extended Data Fig. 2A) and included cytokeratins *KRT8, KRT15*, and *KRT19*, the stem cell marker *CD24*, and the hallmark estrogen response genes *CCND1, TGM2, PRSS23, S100A9*, and *PERP*. Downregulated genes unique to the basal epithelial cells consisted primarily of mitochondrial genes and did not include canonical estrogen response genes. Mature luminal specific downregulated genes included hallmark estrogen response genes *FKBP4, FAM102A, MREG, IGFBP4*, and *SLC9A3R1*. Genes downregulated only in luminal progenitor cells included targets of the ER coregulator GATA3 (*DSTN*, *TM4SF1, MGST12*, and *CLDN7*) and MAPK signaling pathway genes (*HMGN1, PAC1, CDC42, HRAS, PSMB6*, and *RBX1*). Common upregulated genes included *DDIT3* and *EGR1* which have both been shown to suppress breast cancer proliferation^41,42^. Unique genes upregulated with tamoxifen in mature luminal cells included *MGP*, *AREG*, *STC1*, and *HMGS1*, where low expression is associated with endocrine resistance.

To determine differentially regulated pathways with tamoxifen treatment by cell type, we performed enrichment analyses on differentially expressed genes by cell type (Fig 1F, complete list in Supplementary Tables 2-4). Luminal epithelial cells demonstrated strong downregulation of E2-induced and GATA3-induced genes and upregulation of E2-repressed genes, indicating capture of on-target effects of tamoxifen. Basal epithelial cells demonstrated downregulation of E2-induced genes, although less pronounced compared to luminal epithelial cells, and unchanged expression of E2-repressed genes. Of note, fibroblasts did not demonstrate signature enrichment with tamoxifen treatment. Endothelial cells had enrichment of TNFα signaling and induction of estrogen-repressed genes with tamoxifen treatment, indicating a distinct response relative to other cell types. Cumulatively, these findings show that our novel system captures biological effects of tamoxifen treatment in epithelial cells and detects cell populations (i.e., fibroblasts) with differential tamoxifen response. This demonstrates the ability to distinguish heterogeneity in drug response within diverse cell populations in primary human breast tissue.

### Single-cell based response to tamoxifen in T47D cells

Heterogeneity in gene expression and response to treatment occur in distinct subpopulations, even within cell line cultures^43^. To elucidate heterogeneity in response to tamoxifen using our suspension-to-single-cell sequencing platform, we treated the ER+/HER2- breast cancer cell line T47D to recapitulate our human tissue protocol (Fig. 2A). In bulk, T47D has a luminal B gene expression pattern^44^, however, some variability exists at the single cell level. After quality control, processing, and dimensionality reduction, sequenced cells grouped primarily into two subpopulations: groupA (luminal A-like) and a more proliferative groupB (luminal B-like), (Fig. 2B). Cell clustering occurred primarily by cell type rather than treatment condition, which indicates that intrinsic gene expression is the primary driver of clustering (Fig. 2C). As expected, canonical luminal epithelial markers (*KRT8, FOXA1, ESR1*) were uniformly expressed. Proliferation score^40^ was enriched in the groupB subpopulation, Fig. 2E.

**Figure 2.**
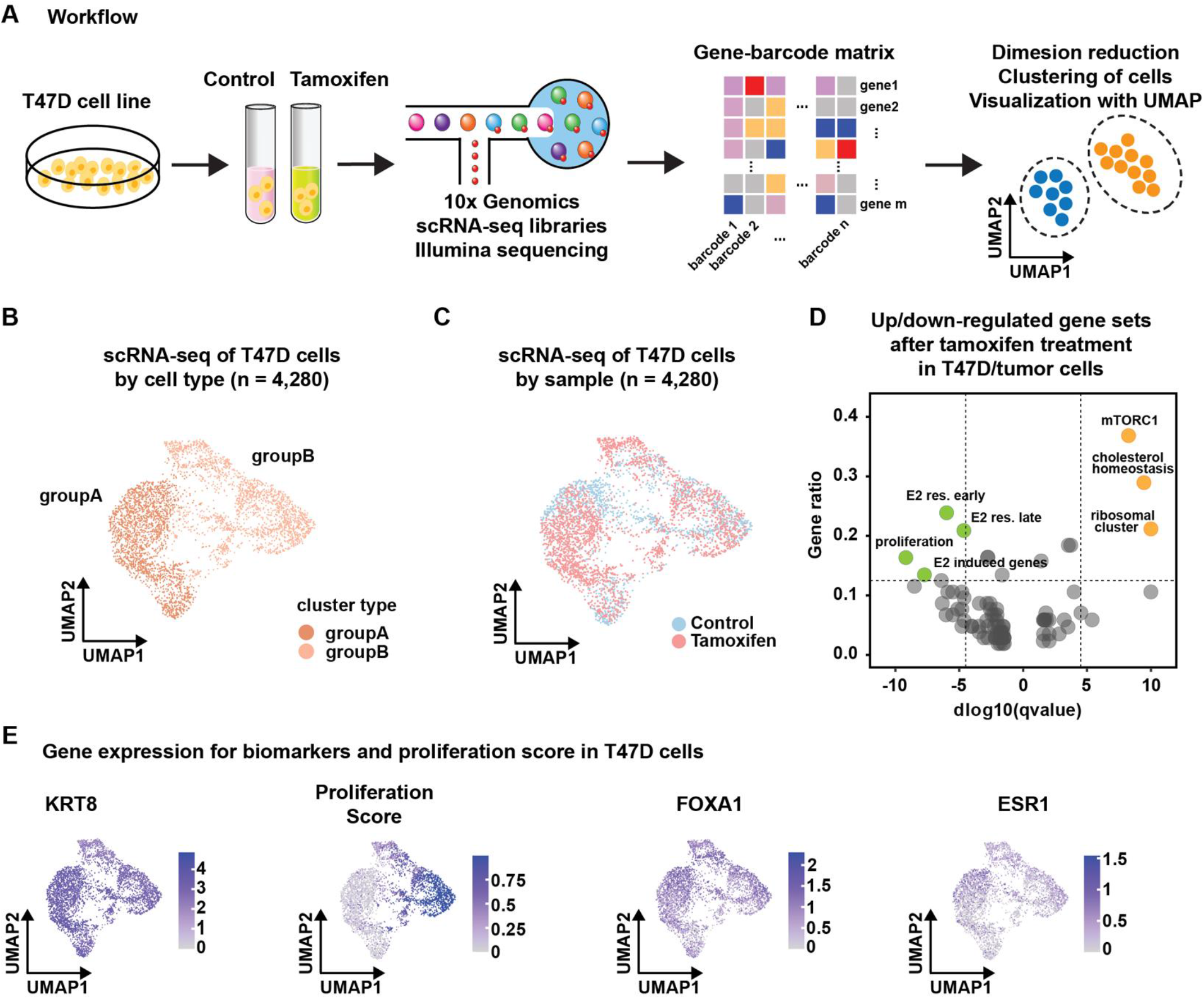
T47D response to tamoxifen at a single cell level. (A) Illustration showing T47D scRNA-seq workflow. (B) UMAP plot of T47D scRNA-seq cells color-coded by groupA (luminal A-like subpopulation) or groupB (luminal B-like subpopulation). (C) UMAP plot of T47D scRNA-seq cells color-coded by treatment condition. (D) Volcano plot showing enriched and depleted gene sets after tamoxifen treatment in T47D cells. Significant gene sets were defined as a gene ratio of > 0.125 and a log10qvalue of < −4.5 or > 4.5, and a thresholding limit of 10 was applied when log10qvalue > 10 for visualization. (E) Feature plots showing the expression of luminal epithelial markers and proliferation score in T47D cells.

Differentially expressed gene analysis was used to determine tamoxifen-regulated genes. In total, we identified 110 downregulated genes and 90 upregulated genes with tamoxifen treatment (complete list in Supplementary Table 5). The downregulated genes in response to tamoxifen treatment were enriched with genes involved in the canonical estrogen-induced pathway and gene signatures indicative of cell proliferation (Fig. 2D). The upregulated genes in response to the treatment were enriched for gene signatures associated with androgen response and activation of the PI3K/mTOR signaling pathway. *DDIT3*, a commonly upregulated gene in normal epithelial cells that can suppress breast tumor growth, was upregulated with tamoxifen in T47D as well. By leveraging 110 downregulated genes (adjusted p-value < 0.01), we established a single-cell signature of tamoxifen response (scTAM-response-T47D).

### Cell type diversity in primary ER+/HER2- human breast tumors

We processed 10 tamoxifen/control pairs from primary ER+/HER2- invasive breast carcinomas using our novel experimental platform. In total, 40,428 cells passed quality control and were annotated using canonical lineage markers (Fig. 3A). Epithelial cells were further classified into malignant (Epi. Tumor) and non-malignant (Epi. Non-tumor) cells by estimating the copy number variation profiles using InferCNV^45^.

**Figure 3.**
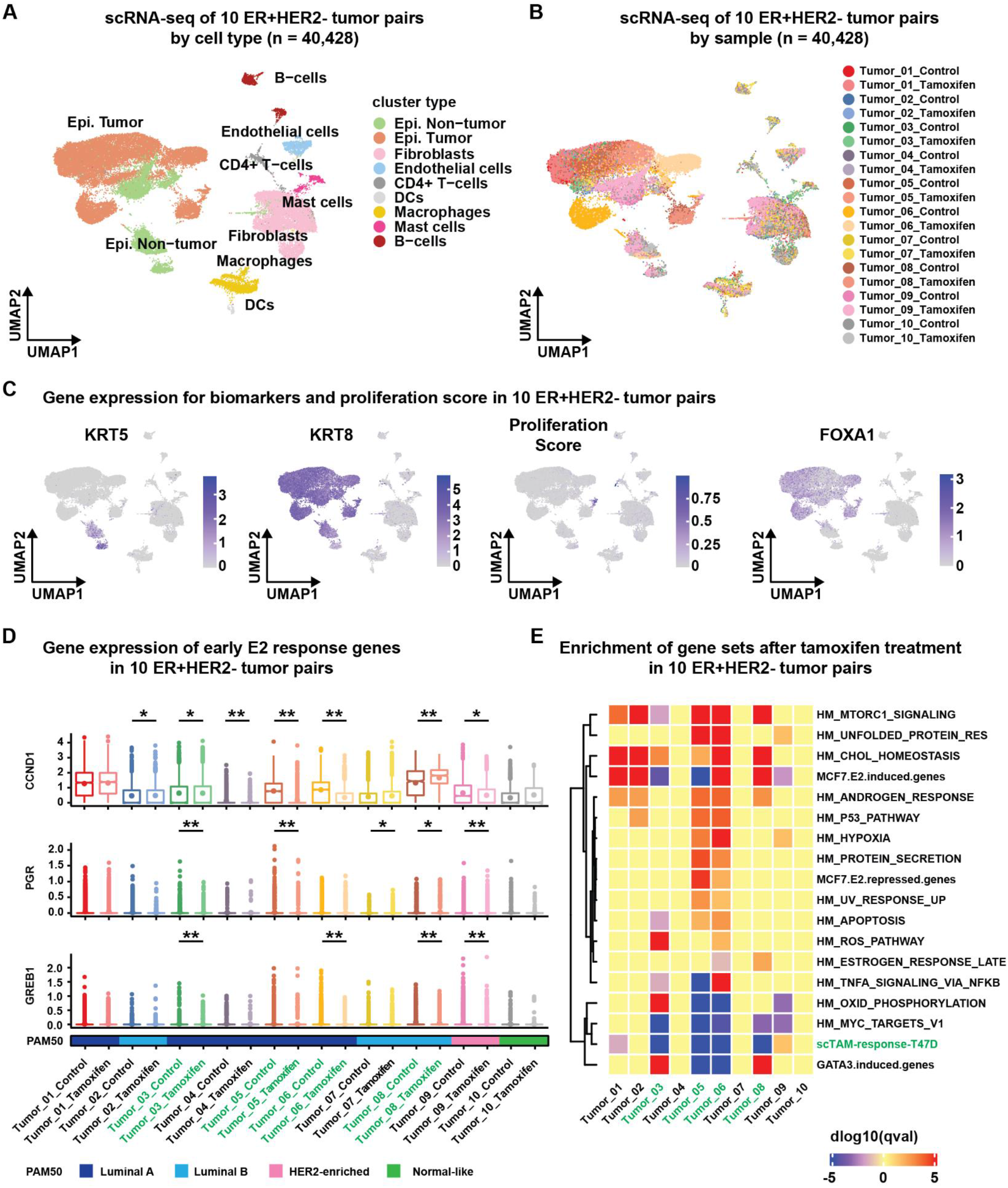
Tamoxifen response in 10 primary ER+/HER2- breast tumors. (A) UMAP plot of all scRNA-seq cells from 10 breast tumor samples. Cells were color-coded by cell type. Malignant epithelial cells (Epi. Tumor) were distinguished from normal epithelial cells (Epi. Non-tumor) by inferred copy number changes using InferCNV. (B) UMAP plot of all cells color-coded by tumor and treatment condition. (C) Feature plots showing the expression of luminal epithelial markers and proliferation score. (D) Box-plots of early estrogen response genes (CCND1, PGR, and GREB1) in the InferCNV+ malignant cells comparing expression in control and tamoxifen treated cells. Significance was determined using the Wilcoxon Sum-Rank test *p-value<0.05, **p-value<0.01. Sample PAM50 subtypes determined on bulk mRNA sequencing are denoted by color-code. (E) Gene set enrichment heatmap showing distinct tamoxifen response within InferCNV+ malignant cells across tumors.

Cells clustered by cell type identity and not by specimen, indicating high quality sequencing. As observed in the normal specimens, there was significant overlap of fibroblasts, endothelial, and immune cells across samples, but not epithelial cells (Fig. 3B). Basal cytokeratin (*KRT5*) expression was present in two distinct populations of non-tumor epithelial cells (Fig. 3C). *FOXA1* expression was enriched in tumor cells compared to non-tumor epithelial and non-epithelial cell types. Despite strong clinical ER-positivity for these tumors and primarily luminal gene expression patterns at both the bulk and single-cell level, *ESR1* transcripts were zero in most tumor cells. This could be due to discordance between transcript abundance and protein levels, or limitations in the depth of sequencing to detect these transcripts.

Cell type abundance and number of sequenced cells was highly variable across samples (Extended Data Fig. 3A). The variability in cell type abundance between samples supports the need for methods to distinguish cell types within heterogeneous populations, which can vary

### Tumor specific changes in estrogen response genes

We leveraged the ability to computationally isolate tumor cells using InferCNV to assess tamoxifen-significantly from specimen to specimen. Single-cell assays can account for distinct cell subpopulations and mitigate effects on downstream analyses. In most specimens, cell type abundance was conserved between control and tamoxifen treatment within samples, with 3 specimens (Tumor_05, Tumor_06, and Tumor_09) showing increased ratio of luminal B/luminal A cells with tamoxifen treatment compared to control.

responsiveness of only malignant cells within tumors. To do this, we evaluated changes in 3 canonical early estrogen response genes (*CCND1, PGR*, and *GREB1*) with tamoxifen treatment for InferCNV+ cells within each tumor (Fig. 3D).

Of the 10 tumor pairs, 6 showed a significant reduction in *CCND1* expression with tamoxifen treatment, although one (Tumor_08) showed increased *CCND1* expression. Five tumors had a significant reduction in *PGR* (encoding the progesterone receptor) and 4 tumors had reduced *GREB1* expression. Overall, this indicates reduced expression of estrogen response genes after tamoxifen treatment in most of these 10 tumors. We did not observe a reduction in *Ki-67* expression in any of the tumors, which may be due to insufficient treatment time for changes in Ki-67 to manifest or low abundance of *Ki-67* transcripts sequenced.

### Variability in tamoxifen response between tumors

To determine tumor-specific tamoxifen response genes and to assess response genes between tumors, we compared differentially expressed genes in malignant cells (i.e. InferCNV+) treated with tamoxifen compared to controls for each tumor (Extended Data Fig. 3B). Upregulated and downregulated genes were highly variable between samples. In general, samples with more downregulated genes had more upregulated genes, which could be due to intrinsic responsiveness or the number of malignant cells sequenced and power to detect gene expression differences.

To assess variability in tumor-specific biological processes regulated by tamoxifen between patients, we assessed enrichment of gene sets. With tamoxifen treatment, estrogen-induced genes^24^ were depleted in 3 tumors (Tumor_03, Tumor_05, and Tumor_09) but enriched in 4 tumors (Tumor_01, Tumor_02, Tumor_06, and Tumor_08) (Fig. 3E). The upregulated genes in response to tamoxifen were enriched in gene sets related to the androgen response, p53, mTORC1 signaling, hypoxia, and apoptosis. MYC target genes and GATA3-induced genes were depleted in several tumors with tamoxifen treatment. The gene set for TNFα signaling was depleted in Tumor_05 and enriched in Tumor_06, highlighting the variability in tamoxifen-mediated inflammatory signaling. To account for variability in sequencing between bulk platforms and our single-cell system, we assessed the previously defined scTAM-response-T47D signature which was depleted in 4 tumors. scTAM-response-T47D genes were enriched in Tumor_09 which was classified as a HER2-enriched bulk PAM50 subtype (Fig. 3E).

### Targeted analysis of tamoxifen-responsive tumors

To determine heterogeneity in response to tamoxifen treatment within specimens, we focused on cells obtained from the 4 tumors with depleted scTAM-response-T47D signature. Cells from these 4 tumors were clustered to visualize cell populations (Fig 4A). Non-epithelial cells showed overlap, whereas epithelial cells clustered by sample (Fig 4B). In InferCNV+ malignant cells, top tamoxifen-regulated genes were plotted (Fig 4C, complete list in Supplementary Table 6). Tamoxifen treatment was associated with decreased expression of estrogen-induced and GATA3-induced genes (Fig 4D), including ER-regulated genes^46^ *KRT7, CCND1, KRT8*, and *XBP1*. Proliferative signatures were depleted in tamoxifen-treated malignant cells. Downregulation of MYC targets, including *FKBP4* and *RANBP1* which alter ER regulated genes, was observed with tamoxifen treatment^43,47,48^.

**Figure 4.**
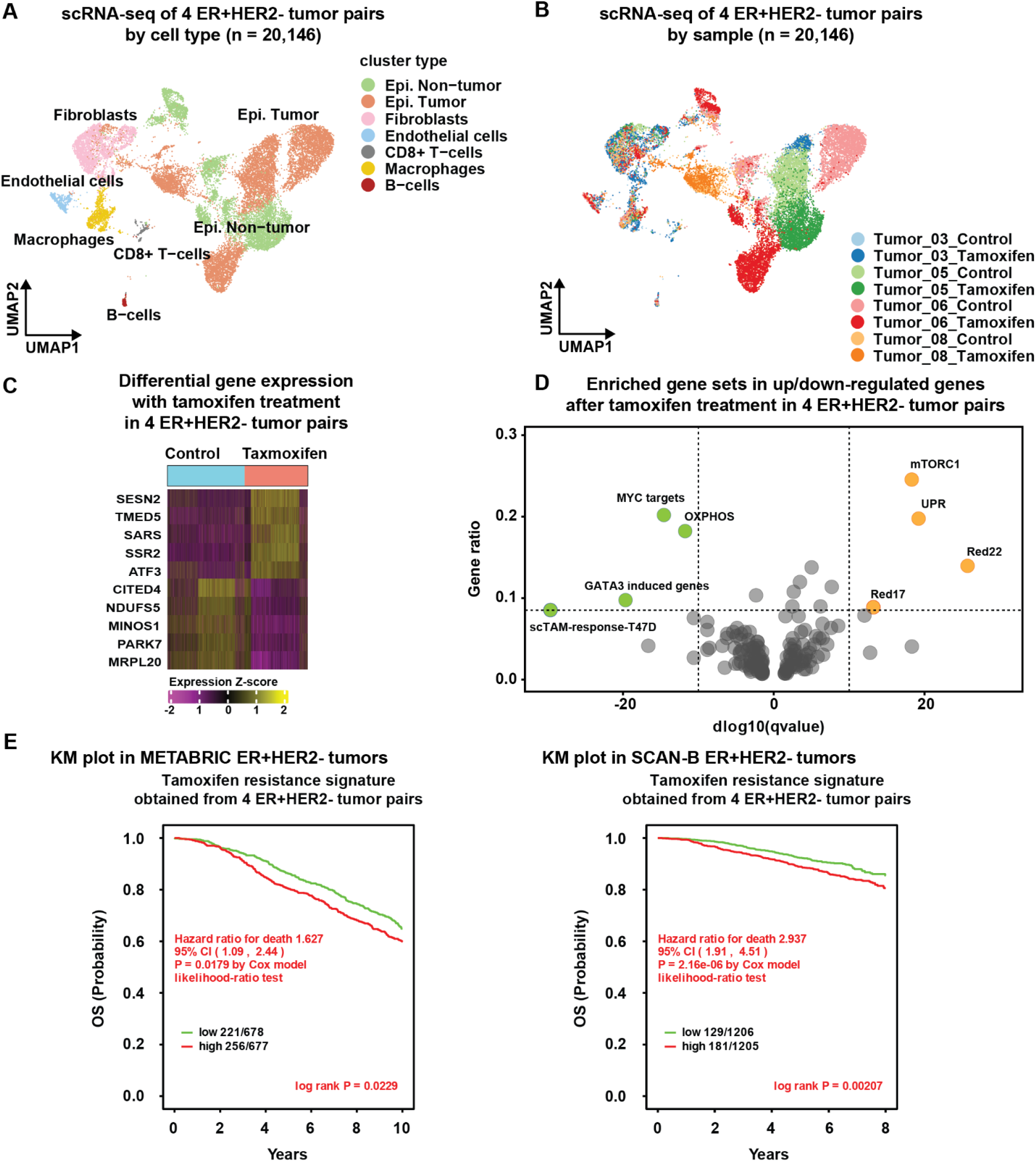
Targeted analysis of 4 tamoxifen responsive tumor pairs. (A) UMAP plot of scRNA-seq cells from 4 tumors that demonstrated depletion of scTAM-response-T47D signature. Cells are color-coded by cell type. (B) UMAP plot of scRNA-seq cells from 4 tumor pairs, color-coded by tumor and treatment condition. (C) Heatmap of upregulated and downregulated gene sets in tamoxifen-treated tumor cells relative to control cells. (D) Volcano plot showing enriched and depleted gene sets after tamoxifen treatment in malignant cells from 4 ER+HER2- tumor pairs. Significant gene sets were defined as a gene ratio of > 0.085 and a log10qvalue of < −10 or > 10 for visualization. (E) Kaplan-Meier (KM) curve for overall survival using two independent clinically annotated datasets with transcriptional data. ER+ patients were assigned a centroid score of our malignant cell-specific tamoxifen resistance signature (scTAM-resistance-M) and stratified into high and low signature score. High signature score is associated with significantly worse overall survival in patients in METABRIC (HR 1.63, p=0.023) and SCAN-B (HR 2.94, p=0.002). Statistical significance was assessed by the log-rank test and the estimates of survival probabilities and cumulative hazard with a univariate Cox proportional hazards model.

Enrichment of several important signatures was also observed among only tumor cells with tamoxifen treatment (Fig 4D, complete list in Supplementary Table 6). Signatures of EGFR and RAS^49^ signaling were enriched including MAPK signaling genes (*MAPK6, VEGFA, DUSP1*, and *DUSP10*). The RAS-RAF-MAPK signaling pathway is known to promote resistance to endocrine therapy^22,50,51^. PI3K/mTORC1 signaling^49^, a targetable pathway in patients with endocrine-resistant tumors^52^, was also enriched in tamoxifen-treated malignant cells. Genes associated with epithelial-mesenchymal transition (EMT)^53^ and FOS/JUN signaling were also enriched and are associated with worse outcomes and poor response to endocrine therapy in breast cancer. The transcription factor *ATF3* involved in FOS/JUN signaling and associated with poor response to endocrine therapy^54^ was also upregulated. Cumulatively, these enriched pathways show rapid adaptive upregulation of genes that may be driving a complex early transcriptional program to promote resistance to endocrine therapy.

To determine variability in tamoxifen response by cell type across specimens, tamoxifen-regulated genes were identified in non-malignant cell populations (Supplementary Table 7). In contrast to malignant cells, fewer differentially regulated genes were seen in fibroblasts and macrophages. Similar to malignant cells, RAS associated genes^49^ were enriched, but this was primarily due to enrichment of mitochondrial genes (*MT2A, MT1X, MT1E*). Tamoxifen downregulated gene sets in malignant cells and fibroblasts did not overlap. Macrophages demonstrated downregulation of hallmark inflammatory response with tamoxifen. Overlap analysis showed that the significant majority of both up and down regulated genes were unique to specific cell types (Extended Data Fig. 4A/B).

These results demonstrate that distinct cell types within primary ER+ breast tumors have unique response to tamoxifen. To assess if a single-cell signature of tamoxifen resistance derived from only malignant cells (scTAM-resistance-M) (Supplementary Table 6) was prognostic in ER+ breast cancer, gene expression data annotated with survival data from METABRIC^55^ and SCAN-B^56^ for patients with ER+/HER2- breast cancer was obtained. Patients were stratified by gene set expression levels. Tumors were ranked according to a signature score based on median expression of the signature genes. Tumors were stratified using median signature score cutoffs into “high” or “low” groups. The impact of signature scores on survival was then assessed with Kaplan-Meier curves. In METABRIC patients, overall survival was significantly worse in patients with tumors that had high expression of the malignant-cell specific tamoxifen response signature (HR 1.63, p=0.023) (Fig 4E). Higher scTAM-resistance-M signature score was also associated with worse survival (HR 2.94, p=0.002) in SCAN-B patients.

### Characterization of resistant malignant cell subpopulations

To measure heterogeneity in tamoxifen response at the single cell level, we developed a response score for individual cells based on the scTAM-response-T47D signature. Each cell was assigned a score as the difference between the treated tumor cell’s centroid signature value and the median centroid value for the matched, untreated tumor cells from the same patient. This metric measures the deviation from the starting state in tamoxifen response genes with tamoxifen treatment and can be quantified for each cell. Differential response to tamoxifen was thus identified between clusters (Fig. 5A). Cluster 2, comprised primarily of LumA-like cells from Tumor_06, displayed a significant reduction in scTAM-response-T47D score, indicating response to treatment. In contrast, clusters 3, 12, and 19 demonstrated increased scTAM-response-T47D scores, indicating unresponsiveness or resistance to tamoxifen treatment. Cluster 19 was the only luminal B (LumB) cluster and the only cluster of cells from multiple tumors (Fig. 5B/C). Although they did not constitute the majority of tumor cells from any tumor, subpopulations that were unresponsive to tamoxifen were detected in 3 of 4 tumors that were responsive when all tumor cells were analyzed jointly.

**Figure 5.**
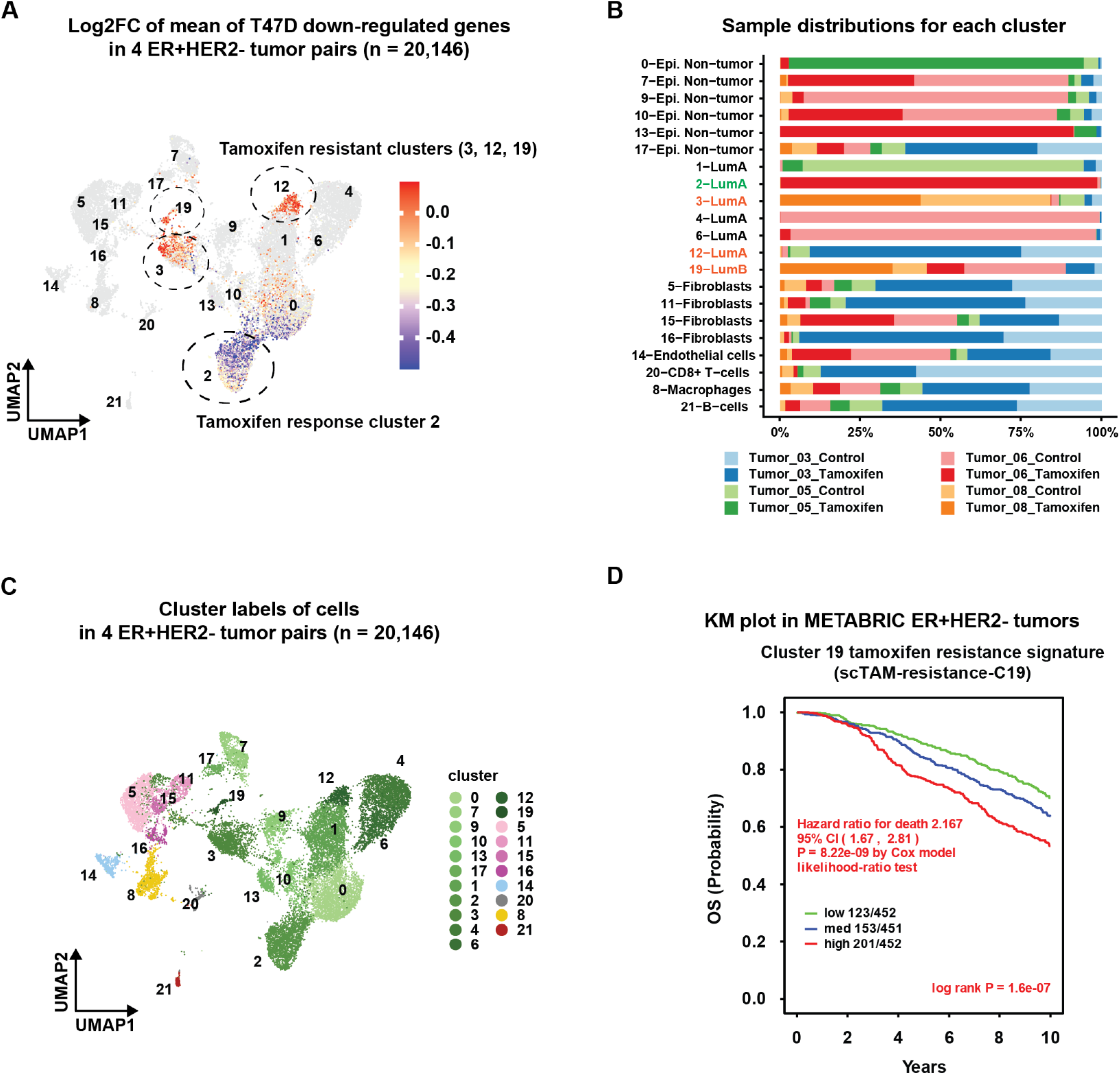
Identification and characterization of resistant tumor cell subpopulations. (A) Individual cells were assigned a score based from the T47D tamoxifen response signature compared to cluster matched untreated signature score. (A) Application of response score to UMAP plot demonstrated 3 distinct clusters with enriched signature score on treatment (Cluster 3, 12, 19) and one tamoxifen sensitive cluster with depletion of response score (Cluster 2). (B) Stacked bar chart showing distribution of cells within 22 distinct clusters, color-coded by tumor and treatment condition. (C) UMAP plot of scRNA-seq cells from 4 tumors with depleted T47D tamoxifen response score. Cells were color-coded by cluster. (D) Kaplan-Meier (KM) curve demonstrates the prognostic significance of the cluster 19 signature (scTAM-resistance-C19), which was evaluated using the METABRIC dataset. Higher signature score predicted worse overall survival (HR 2.17, p<0.001). Survival curve differences were calculated by the log-rank test and the estimates of survival probabilities and cumulative hazard with a univariate Cox proportional hazards model.

To determine features of these unresponsive subpopulations, we identified differentially expressed genes for each of the 3 clusters compared to the responsive cluster 2 (Extended Data Fig. 5A, complete list in Supplementary Tables 8-10). Clusters 3 and 12 were both characterized by enrichment of RAS signaling pathways and EMT, both of which can contribute to reduced estrogen responsiveness. Cluster 19 was characterized by high Ki-67 and enrichment of proliferative signatures^57–59^ (Extended Data Fig. 5B). Highly proliferative cells could indicate a population that was dividing or committed to dividing during the short-term treatment, thus completing the cell cycle under treatment rather than being truly “unresponsive”. Several reports have demonstrated that high Ki-67 is associated with worse outcome in endocrine therapy-treated patients and that a reduction in Ki-67 is a key clinical metric of response to endocrine therapy^27,28,60^. Interestingly, all 3 clusters had increased expression of *TACSTD2* encoding TROP2, the target of the antibody-drug conjugate Sacituzumab govitecan which has proven efficacy in metastatic ER+/HER2- breast cancer^61^. Therefore, all 3 clusters identified as resistant were characterized by gene expression patterns of known mechanisms of resistance to endocrine therapy, and have increased expression of a druggable gene that could be used to deliver precision, subpopulation specific therapy. These findings support the validity of this model to identify true differential response within tumor cell subpopulations and identify potential treatment strategies.

To assess whether resistant subpopulations contribute to poor outcomes, we used clinically annotated transcriptional data from ER+/HER2- tumors in METABRIC and SCAN-B. Resistant cluster signatures were generated for each of the 3 clusters; scTAM-resistance-C3, scTAM-resistance-C12, and scTAM-resistance-C19 (Supplementary Tables 8-10). Tumors were ranked by median and tertile cutoffs and survival assessed by Kaplan-Meier curves. Tumors were stratified using tertile signature score cutoffs into “high”, “med”, or “low” groups. The scTAM-resistance-C19 signature demonstrated the most robust stratification for overall survival (Fig. 5D), with high score corresponding to reduced survival in METABRIC (HR 2.2, p<0.001) and in SCAN-B (HR 1.9, p=0.002). High expression of the scTAM-resistance-C3 signature was significantly associated with reduced overall survival in SCAN-B (HR 2.2, p=0.03) but not in METABRIC. Similarly, high expression of the scTAM-resistance-C12 signature was associated with reduced survival in SCAN-B (HR 1.9, p=0.003) but not in METABRIC (Extended Data Fig. 6A).

Lastly, to determine if these resistant clusters were enriched in endocrine-resistant tumors, bulk RNA-seq data was obtained from patients with unresectable or metastatic ER+/HER2- breast cancer prospectively treated with endocrine therapy^51^. Serial biopsies were taken and samples were annotated as “sensitive” or “resistant” based on clinical response. Resistant tumors had significantly higher expression for all three resistant signatures relative to sensitive tumors (−0.11 vs 0.16 for cluster 3, p=0.00037; - 0.12 vs. 0.21 for cluster 12, p=0.00081; −0.12 vs. 0.26 for cluster 19, p=0.00031) (Extended Data Fig. 6B).

## DISCUSSION

Assessing heterogeneity in response to therapy within primary human solid organ tumors represents a significant challenge. For the first time, we report the use of single-cell transcriptional profiling coupled to a primary human tumor ex vivo experimental platform to dissect drug response within and across tumors. Using this platform, this study identified and characterized distinct responses to tamoxifen within normal human breast tissue and primary ER+/HER2- human breast tumors. The ability to successfully distinguish differential response has several important implications. First, it allows for computational analysis of distinct cell populations, such as malignant cells, which can be used to derive more precise signatures of response. Herein, we generated signatures from computationally-identified tumor cells and showed that these were highly prognostic in patients with ER+ breast tumors across multiple large datasets. This could be especially important in lobular breast cancers in which malignant cells are significantly outnumbered by other cell types^29^ as well as other solid organ tumors with variable malignant and stromal compositions. These findings demonstrate that dissecting the response to tamoxifen in specific cell compartments has the potential to improve precision in understanding therapeutic response. This platform could thus be used to uncover mechanisms that are potentially masked in bulk methodology.

We have additionally shown the ability to assess differential sensitivity to therapy within malignant cells of the same tumor. In some tumors, resistance to therapy may be driven by low-abundance cell populations that are masked in bulk sequencing and require single-cell methods to dissect. Importantly, we present a single cell-derived response score that identified 3 low-abundance cell populations that are unresponsive to tamoxifen. These resistant cells were identified within tumors that would have been classified as responsive (depleted E2-induced genes) at the tumor level. Thus, the ability to distinguish these cell populations may unmask hidden resistant cell populations. Supporting the accuracy of this characterization, all 3 populations had features of described mechanisms of resistance to endocrine therapy and were enriched in endocrine resistant tumors. Interestingly, these subpopulations all had increased expression of TROP2, the target of the antibody-drug conjugate Sacituzumab govitecan. The presence these low abundance tamoxifen resistant cells could identify patients in whom a precision medicine dual treatment strategy of endocrine therapy for the sensitive subpopulation and Sacituzumab govitecan for the TROP2-expressing resistant subpopulation is more efficacious than endocrine therapy alone. Further study is needed to develop clinically actionable strategies based on precise characterization of tumor cell subpopulation response to therapy.

We, and others, have previously studied MAPK signaling, which causes reduced sensitivity to endocrine therapy in ER+ breast cancer^22,62–65^. We identify both a MAPK-mediated adaptive response to tamoxifen in tumor cells and a tamoxifen-resistant subpopulation characterized by enrichment MAPK signatures. We also show in primary human tumors that tamoxifen treatment results in upregulation of *DUSP1* and *DUSP10*, which encode inhibitory phosphatases, despite their depletion in the inherent-resistance clusters. This indicates that induction of *DUSP1* and *DUSP10* with tamoxifen may be an important mechanism to prevent the bypass of estrogen signaling via upregulation of MAPK activity. Further studies of protein activity are needed to dissect precise adaptation of ERK/MAPK signaling components in response to tamoxifen and its contribution to endocrine therapy resistance.

We recognize that our study has several limitations. We present a small sample size with limited power for subgroup analysis based on tumor or patient-specific factors such as menopausal status and ductal/lobular histology. Additionally, the analysis of resistant cell populations was based on cells from tumors that were determined to be sensitive to tamoxifen as measured by depletion of a platform-matched T47D derived signature, which may limit the ability to characterize resistant populations. Nevertheless, this work represents the first model system to annotate distinct treatment responses within individual cells of primary human breast tumors and offer a treatment target for resistant cells.

In conclusion, we present a novel ex-vivo drug treatment platform coupled to single-cell RNA sequencing for human breast tumors. Through this platform, we identified distinctions in drug response of cell populations in normal human breast tissue and primary human tumors. Several mechanisms of resistance to tamoxifen were identified in resistant malignant cell subpopulations within human tumors, and signatures of those subpopulations predicted poor outcomes in patients with ER+ breast tumors. Further studies are needed to determine clinically actionable strategies to identify patients at risk for therapeutic failure based on the presence of low-abundance resistant subpopulations and determine co-treatment strategies to overcome resistance in these patients.

## METHODS

### Patient specimens

Patients were identified in the breast cancer surgery clinic with a diagnosis of ER+/HER2-negative breast cancer or in the plastic surgery clinic as patients without breast cancer undergoing reduction mammaplasty. Breast tissue was obtained from patients who did not receive any preoperative therapy. Written informed consent was obtained from all patients under a protocol that was approved by the Institutional Review Board (IRB) at the University of North Carolina at Chapel Hill. Consent included access to de-identified patient data which was obtained through an honest broker. Participants were not compensated for participation.

### Tissue processing and dissociation

Primary tissue specimens were obtained in the operating room suite and placed in DMEM/F12 (Gibco) media with 1% Penicillin-Streptomycin (Life Technologies). Specimens were transferred immediately to the laboratory where the specimens were processed into single cell suspensions by sharply mincing into 2-4 mm fragments. Enzymatic dissociation was then performed using Gentle Collagenase/Hyaluronidase (Stemcell Technologies Inc. 07919) in DMEM/F12 supplemented with 5% BSA (Sigma Aldrich), Hydrocortisone (Stemcell Technologies Inc.), HEPES (Corning), and Glutamax (Gibco) for 16 hours at 37°C with cell agitation. The cells were gently centrifuged and washed twice with PBS (Gibco) supplemented with Fetal Bovine Serum (FBS) and HEPES buffer. Cells were resuspended in cold ammonium chloride solution (Stemcell Technologies Inc. 07800) and incubated at room temperature to remove red blood cells. Cells were centrifuged and briefly trypsinized in warm 0.05% Trypsin-EDTA (Gibco) and DNase I (Stemcell Technologies Inc. 07900). Cells were further dissociated with dispase (Stemcell Technologies Inc. 07923) and DNase I. Cells were centrifuged and washed then resuspended in DMEM/F12 with 10% FBS and 1% Penicillin-Streptomycin. Cells were then strained using multiple rounds of sequential straining with 40 μm mini strainers (Pluriselect USA Inc.) to remove cell debris.

### Tamoxifen suspension

Cells were counted using fresh trypan blue (Invitrogen) and the Countess II cell counter (Life Technologies). Dissociated single cells (n=100,000) were placed in suspension on an Ultra-Low attachment microplate (Corning 3474) in DMEM/F12 media supplemented with 10nM estradiol (Sigma Aldrich). Paired suspensions were treated with control media or media containing 10 μM/L 5-OH Tamoxifen. Suspensions were incubated at 37°C with 5% CO2 for 12 hours after which cell viability was assessed with trypan blue. Graphical representation of workflow from tumor resection to data analysis is shown in Fig. 1A.

### Cell line culture

We obtained the ER+/HER2- cell line T47D from the American Type Culture Collection (ATCC). T47D cultures were maintained at 37°C with 5% CO2 in DMEM with 10% FBS and 1% Penicillin-Streptomycin. Cells were trypsinized and suspended in control or tamoxifen containing media on an Ultra-Low attachment microplate for 12 hours, assessed for viability, and processed for single cell RNA sequencing libraries.

### Single-cell RNA-seq

Single-cell RNA sequencing was done with the 10x Genomics Chromium Next GEM Single Cell 3’ v3.1 reagent kits according to the manufacturer’s revision D user guide. A total of 7,000 to 10,000 viable cells were targeted per library. Libraries were sequenced on the NextSeq500 (Illumina) and NextSeq2000 (Illumina) platforms with pair-end sequencing and single indexing according to recommended manufacture protocols.

### Bulk RNA sequencing and analysis

Tumor specimens were obtained from clinical formalin-fixed, paraffin-embedded (FFPE) blocks. Slides were reviewed for blocks with high tumor concentrations and unstained slides cut. RNA was isolated from the FFPE tissue by use of the High Pure FFPE RNA Micro Kit (Roche 04823125001). Slides were cut into 2 10 micron sections, and were incubated in Hemo-De (Scientific Safety Solvents HD-150) and then washed in 100% and 70% ethanol. The tissue was then air-dried at 55°C. Tissue Lysis Buffer and Proteinase K were then added to the tissue pellet and incubated at 55°C. Binding Buffer and ethanol were then added to the tissue pellet and the supernatant was filtered out via filter tube. DNase Solution and DNase Incubation Buffer were then added to the filter tube and incubated at room temperature. A series of washes were then performed using Wash Buffer I and Wash Buffer II. Elution Buffer was then added to the filter tube, incubated at room temperature, and RNA was then eluted. The purified RNA concentration was then determined via nano-drop.

RNA-seq reads were aligned to the human genome hg38 (gencode_v36) from Genomic Data Commons using STAR v2.7.6a. Transcripts were quantified using Salmon v1.4.0. The matrices were upper quartile fixed normalized (UQN) where quantile (0.75) = 1000.

### PAM50 subtype classification in bulk RNA-seq data

The UQN data was log2 transformed and, after assessing for batch effect, no differences were observed between the two datasets. For PAM50 subtype classification, we applied a HER2/ER subgroup-specific gene-centering method as described in the supplemental methods of Fernandez-Martinez et al.^66^ which requires the log2-transformed UQN data and the IHC status for all samples assayed by RNA-seq. The gene expression values of the PAM50 genes were then HER2/ER-subgroup-specific gene-centered, and the PAM50 predictor^67^ was applied using the provided centroids to assign subtype calls using correlation values for all primary tumors.

### Determining Ki-67 on matched clinical specimens

Slides from clinical FFPE blocks were obtained from the clinical pathology specimens for all tumors. Immunohistochemistry (IHC) for Ki-67 antigen was performed on formalin-fixed, paraffin-embedded tissue sectioned at 4 microns. Staining was performed using the Leica Bond III Autostainer system. Slides were dewaxed in Bond Dewax solution (AR9222) and hydrated in Bond Wash solution (AR9590). Heat induced antigen retrieval was performed at 100°C in Bond-Epitope Retrieval solution 1 pH-6.0 (AR9961). After pretreatment, slides were incubated with Ki-67 Antibody (MIB-1, Dako) at 1:100 for 30 minutes followed with Novolink Polymer (RE7260-K) secondary. Antibody detection with 3,3’-diaminobenzidine (DAB) was performed using the Bond Intense R detection system (DS9263). Stained slides were dehydrated and cover slipped with Cytoseal 60 (8310-4, Thermo Fisher Scientific). A positive control tissue was included in the run. For future reference, IHC slides were digitally imaged in the Aperio AT2 (Leica) using 20x objective. Ki-67 was scored on the glass slide of whole tissue sections by a breast pathologist (BC) according to the Ki-67 IHC MIB-1 pharmDx (Dako Omnis) Interpretation Manual for Breast Carcinoma. The Ki67 pharmDx score (%) was calculated as number of Ki-67 staining viable invasive tumor cells divided by the total number of viable invasive tumor cells, multiplied by 100. A minimum of 2000 cells were scored for each slide. Slides were read by a breast cancer clinical pathologist (BCC).

### Data processing and cluster annotations

The filtered feature barcode matrices were generated by Cell Ranger from 10x Genomics for our scRNA-seq dataset built from normal breast tissue specimens, the T47D cell line, and ER+ breast cancer tumor specimens with or without tamoxifen treatment. Data processing and subsequent downstream analyses were divided into four analysis blocks, namely ABlock1-4. In ABlock1, two paired normal samples were analyzed by processing two normal breast tissue specimens into tamoxifen/control pairs, producing two paired samples (four samples in total). ABlock2 was devoted to the study of T47D cells. ABlock3 focused on 10 pairs of ER+ breast cancer tumor samples. Finally, ABlock4 concentrated on targeted analysis of four pairs of ER+ breast cancer tumors that were selected based on response to tamoxifen measured by depletion of scTAM-rsponse-T47D signature. For each analysis block, Seurat objects were built from the filtered feature barcode matrix for samples assigned to the analysis block using the Seurat R package^68,69^.

Outlier cells were defined by three metrics: low number of UMI counts, low number of genes expressed, and high percent mitochondrial read count (> 25%). For ABlock1 and ABlock4, we used the following threshold values to identify outliers: the number of UMI counts < 2,000 and the number of genes expressed < 1,000. For ABlock2 and ABlock3, we used more stringent threshold values using UMI counts < 5,000 and the number of genes expressed < 2,000 to select cells with higher quality. The purpose of ABlock2 was defining the upregulated gene signatures due to tamoxifen treatment. We defined this gene signature with cells of higher quality to be optimally representative. The purpose of ABlock3 was selecting reliable tamoxifen response pairs among 10 pairs of ER+ breast cancer tumors. We used higher thresholding to select higher quality cells to exclude specimens with lower number epithelial cells of higher quality. The outlier cells according to these criteria were removed before downstream analyses.

After removing doublets by DoubletFinder^70^ to make the final Seurat object for each sample, we applied Seurat’s *merge(*) function^71^ to combine multiple Seurat objects from samples associated with an analysis block for each analysis block. The log normalized procedure was applied to the gene expression matrix. Subsequently, a scaling operation was performed on the expression values of the 2,000 most variably expressed genes to facilitate subsequent principal component analysis (PCA). The confounding effect of the percentage of mitochondrial genes^71^ was subsequently regressed out. A reduction in dimensionality was performed through the summarization of the 2,000 most variably expressed genes into 50 principal components (PCs) via PCA. The cells were then projected onto a uniform manifold approximation and projection (UMAP) embedding via Seurat^69^ and ggplot2^72^ R packages.

For further clustering analysis, a shared nearest-neighbor graph was constructed using 30 PCs, and Louvain clustering was performed with a resolution of 0.8.

### Cell type annotation

The cell type annotations were performed using the SingleR^73^ R package, which leverages reference transcriptomic datasets of well-defined cell types. The normalized expression values obtained from human bulk RNA-seq data sourced from Blueprint and ENCODE were utilized as reference datasets with the SingleR^73^ framework. The reference dataset is available via the celldex R package 73.

To facilitate analysis for each analysis block, a merged cohort dataset was created by integrating the raw count matrices of samples assigned to each analysis block. Subsequently, a canonical correlation analysis (CCA) was performed using Seurat^71^ to correct for batch effects, and cell clustering was performed utilizing the Louvain clustering method. The representative cell-type for each cluster in the merged dataset was annotated with a cell type label based on the predominant cell type within each cluster.

### Distinguishing malignant cells from non-malignant epithelial cells

To differentiate tumor cells from normal epithelial cells, copy number events for each cell cluster were estimated using the InferCNV R package^45,74^ utilizing a normal background comprised of immune cells and endothelial cells. The cell clusters were specified in the InferCNV annotations file, enabling the estimation of CNVs at the level of these clusters. The epithelial cells were subsequently stratified into epithelial tumor and epithelial normal cells by plotting scatter plots of the CNV values and the correlation values with the top 5% of cells with high CNV values^40,75^. The tumor epithelial cells were used for downstream analyses for characterizing the molecular subtypes of tumor cells and identifying molecular signatures of tamoxifen resistance.

### PAM50 subtypes and proliferation score on single cells

Molecular subtype estimation was performed by the nearest centroid method using four-centroids of LumA, LumB, HER2-enriched, and Basal, obtained from scRNA-seq profiles and the molecular subtype classifications of single cells by SCSubtype^40^ of human breast cancers. The proliferation score was computed by the averaged normalized expression of 11 genes (*BIRC5, CCNB1, CDC20, NUF2, CEP55, NDC80, MKI67, PTTG1, RRM2, TYMS* and *UBE2C*)^40^.

### Differential gene expression and pathway enrichment analysis

The differential gene expression between groups was determined using Seurat’s *FindMarkers(*) function^71^, with log2FC (log2 fold change) threshold set at 0.25 and minimum percentage of expressed cells set at 0.25. The markers were then further filtered based on an adjusted p-value threshold of less than 0.01. The top 5 upregulated and downregulated genes after tamoxifen treatment in basal, luminal progenitor, and mature luminal epithelial cells were displayed with heatmaps in Extended Data Fig. 1C for visual clarity. The upregulated and downregulated genes in tamoxifen resistant clusters (Cluster 3, 12, 19) compared to the tamoxifen responsive cluster (Cluster 2) were also displayed with heatmaps in Extended Data Fig. 2B. The enrichment of cancer hallmark gene sets^76^ and a curated list of breast cancer relevant gene signatures^59^ were assessed using hypergeometric tests with *enricher(*) function from the clusterProfiler package^77^ with q-value threshold of less than or equal to 0.01.

### Survival analysis of single cell signatures

The clinical significances of gene signatures were estimated by Kaplan-Meier curves. The signature score was defined by the median expression of the signature genes. Tumors in METABRIC dataset^55^ (EGAS00000000083) or SCAN-B dataset^56,78^ (GSE202203) were stratified into high/low (by 50th percentile) and high/med/low (by tertiles) groups by signature scores. The gene expression data and clinical data for METABRIC were download from cBioPortal for cancer genomics^79,80^. Malignant-cell-specific tamoxifen resistance signature (scTAM-resistance-M) was generated from the top 300 downregulated genes ranked by fold change after selecting genes with log2FC > 0.25 and adjusted p-value < 1e-6, in ABlock4 using our four pairs of tumor samples. The Kaplan-Meier plots for this signature can be found in Extended Data Fig. 3. After comparing transcriptomic profiles in tamoxifen resistant clusters (Clusters 3, 12, and 19) with tamoxifen response cluster (Cluster 2), we generated three gene signatures of tamoxifen resistance from genes upregulated in tamoxifen-resistant clusters (see cluster specific supplemental files). Kaplan-Meier curves were plotted using the survival R package for overall survival to demonstrate the impact of stratification by the enrichment of resistant subpopulation signatures (Extended Data Fig. 5A). Statistical significance was assessed using the log-rank test with significance defined as p<0.05.

### Enrichment of resistant clusters in endocrine therapy-resistant tumors

To determine enrichment of resistant cluster signatures in endocrine therapy-resistant tumors, we obtained clinically annotated bulk RNA-seq data from patients prospectively followed while on endocrine therapy^51^. Serial biopsies were obtained and response to endocrine therapy was assessed clinically as “sensitive” or “resistant”. Centroid values for each of the 3 resistant subpopulation signatures were calculated as the median gene expression value of signature genes for each sample. Signature scores were compared between sensitive and resistant samples using violin plots (Extended Data Fig. 5B) and statistical significance was determined using the Wilcoxon Sum-Rank test using the ggpubr R package.

## Supporting information

Supplementary Table 1: Comparison of Normal_01 with/without 12hr suspension

Supplementary Table 2: Comparison of tamoxifen and control treated basal epithelial cells (BEp) from normal samples

Supplementary Table 3: Comparison of tamoxifen-treated and control-treated luminal progenitor cells (LEp_prog) from normal samples

Supplementary Table 4: Comparison of tamoxifen and control treated mature luminal cells (LEp) from normal samples

Supplementary Table 5: T47D response to tamoxifen

Supplementary Table 6: Response to tamoxifen in InferCNV+ malignant cells within the selected 4 pairs of ER+/HER2- tumors

Supplementary Table 7 Comparison of tamoxifen and control treated non-malignant cell subpopulations within the selected 4 pairs of ER+/HER2- tumors

Supplementary Table 8: Comparison of upregulated and downregulated genes in resistant cell subpopulation cluster 3 compared to cluster 2

Supplementary Table 9: Comparison of upregulated and downregulated genes in resistant cell subpopulation cluster 12 compared to cluster 2

Supplementary Table 10: Comparison of upregulated and downregulated genes in resistant cell subpopulation cluster 19 compared to cluster 2

## Data Availability

Raw data (10x FASTQs) and processed data for bulk and single-cell RNA-seq data have been deposited at the National Institutes of Health (NIH) Database of Genotypes and Phenotypes (dbGaP) (https://www.ncbi.nlm.nih.gov/gap/) and is available under the accession number phs003186.v1.p1.

## Code Availability

All original code has been deposited on the Github, and it is publicly available at the Github repository BC_tamoxifen_response (https://github.com/hyunsoo77/BC_tamoxifen_response). Additional information regarding data analysis is available upon request.

## ACKNOWLEDGEMENTS

This work was presented, in part, as a Spotlight Poster Discussion at the 2022 San Antonio Breast Cancer Symposium. Dr Philip Spanheimer is supported by the National Institutes of Health grant P50CA058223 (P.I. CM Perou) and a Junior Faculty Award from the Society of University Surgeons Foundation. This work was supported in part by P30CA016086 UNC Lineberger Comprehensive Cancer Center Core Support Grant and a grant from the NIH/National Cancer Institute (R01-CA273444-01) to H.L.F. We thank Albert Wielgus in the Pathology Services Core for expert technical assistance with Histopathology/Digital Pathology including tissue sectioning, immunohistochemical staining, and imaging. The PSC is supported in part by an NCI Center Core Support Grant (P30CA016086).

## CORRESPONDANCE

Philip M. Spanheimer MD

Lineberger Comprehensive Cancer Center

University of North Carolina at Chapel Hill

170 Manning Drive, Suite 1149 Chapel Hill, NC 27599-7213 USA Phone: 919-966-5221; Fax: 919-966-8806

## EXTENDED DATA

**Extended Data Fig. 1.**
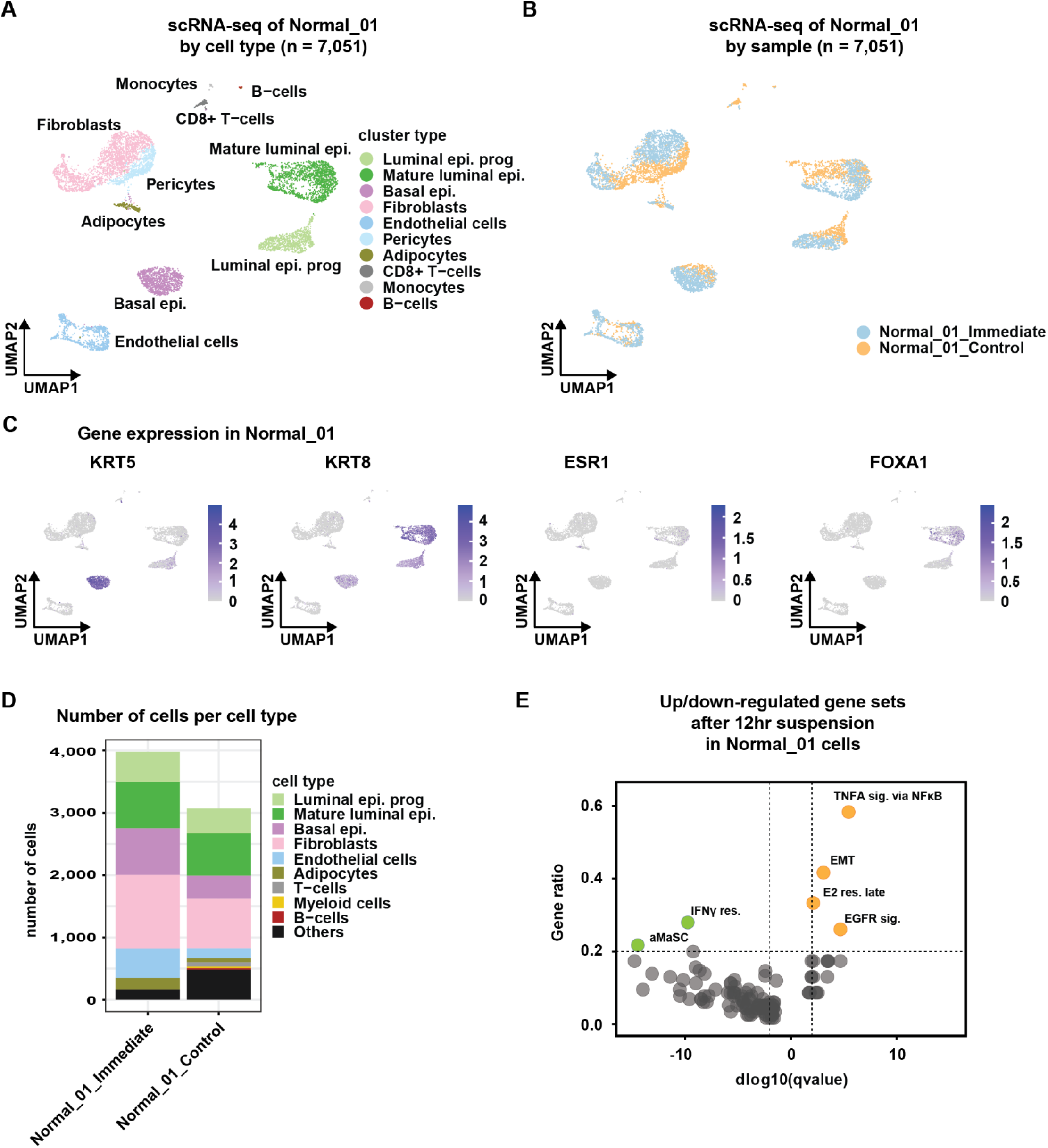
Effect of time in suspension on cell population abundance and gene expression. (A) UMAP plot of all cells from Normal_01 with immediate library creation and control suspension color coded by cell type. (B) UMAP plot of cells color coded by sample and demonstrating clustering by cell type and not by time in suspension. (C) Feature plots of canonical breast epithelial markers. (D) Bar chart comparing the abundance of discrete cell populations in the immediately sequenced sample and after 12 hours in suspension demonstrating fewer cells after time in suspension without systematic depletion of any cell population. (E) Volcano plot comparing gene set enrichment and depletion in up and down regulated genes with time in suspension. Significant gene sets were defined as a gene ratio of ≥ 0.2 and a log10qvalue of < −2 or > 2.

**Extended Data Fig. 2.**
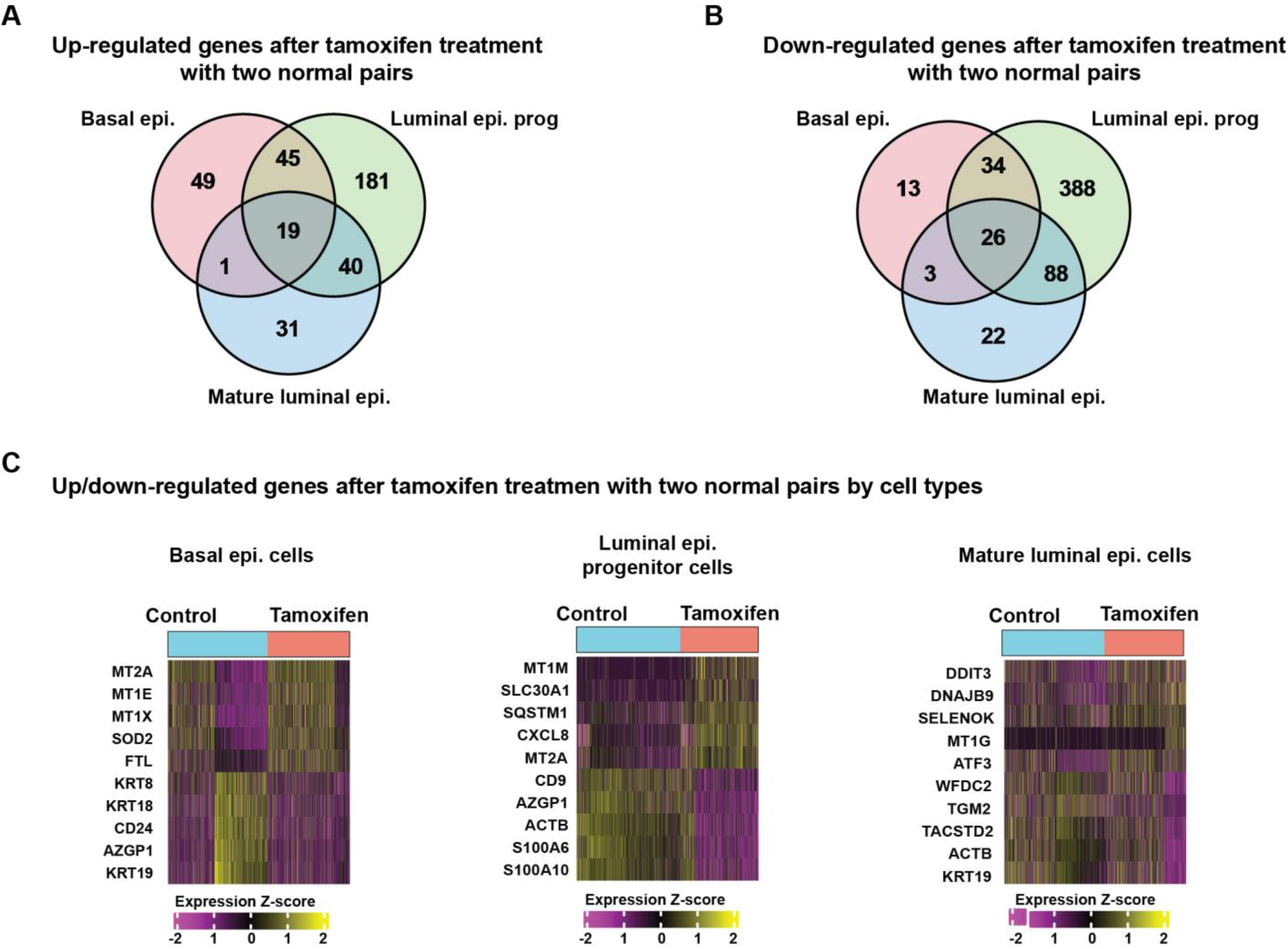
Cell type-specific tamoxifen regulated genes in normal human breast tissue. (A) Venn diagram showing overlapping number of up-regulated genes between basal, luminal, and mature luminal normal breast cells treated with tamoxifen. Significance was defined as q-value < 0.01. (B) Venn diagram showing overlapping number of down-regulated genes between basal, luminal, and mature luminal normal breast cells treated with tamoxifen. (C) Cell type-specific heatmaps of top up- and down-regulated genes by cell type.

**Extended Data Fig. 3.**
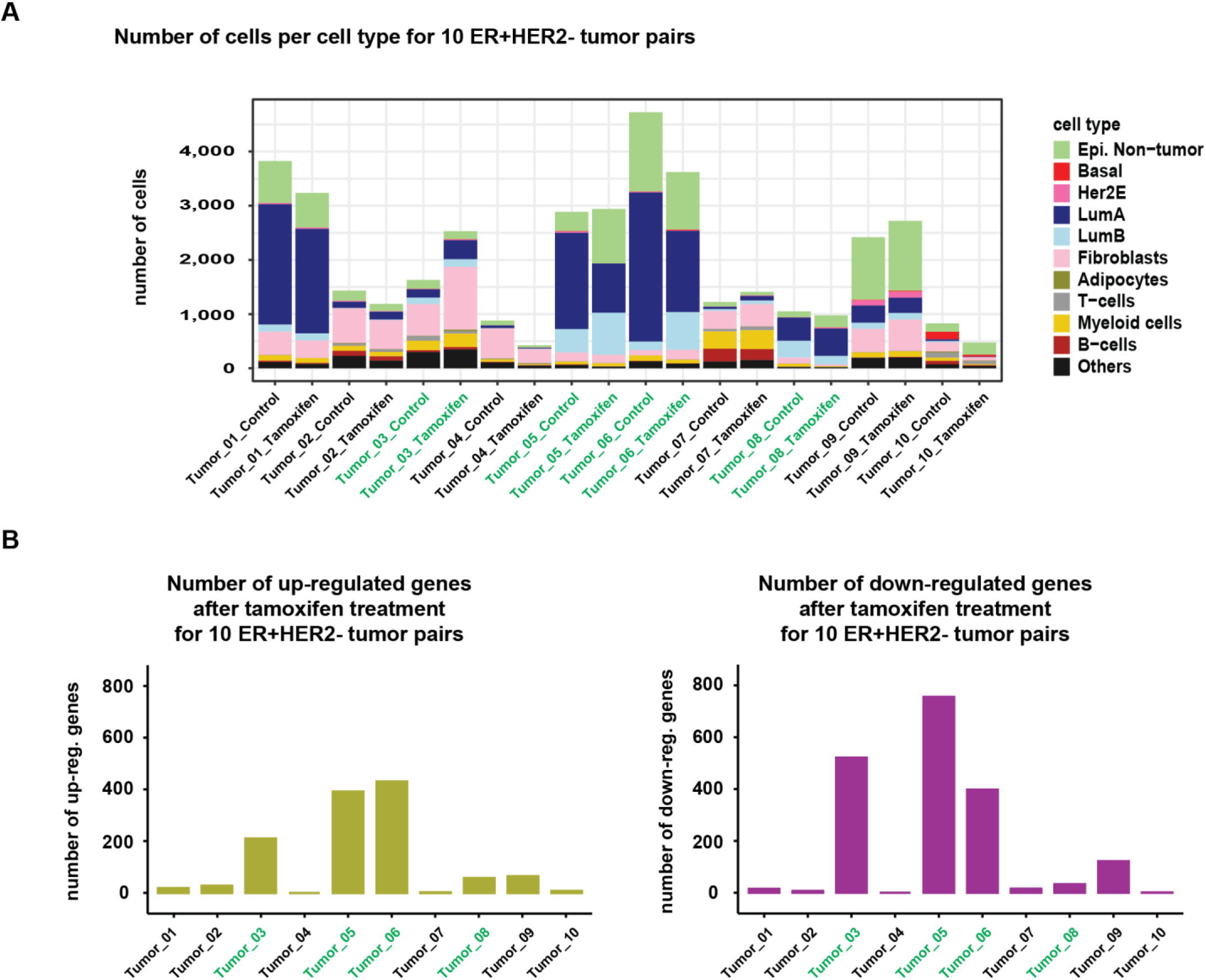
Tumor-specific cell annotations and tamoxifen-regulated genes. (A) Bar chart comparing the total number of cells sequenced per treatment condition, color-coded by cell type. Malignant cells were further classified into PAM50 intrinsic subtypes by the nearest neighbor method using centroid vectors obtained from single cells defined by a single-cell method of intrinsic subtype classification (SCSubtype)^40^. (B) Bar chart showing number of up- and down-regulated (significance defined as log2fold change > 0.25 and q-value < 0.01) with tamoxifen treatment in the computationally identified malignant tumor cells by tumor.

**Extended Data Fig. 4.**
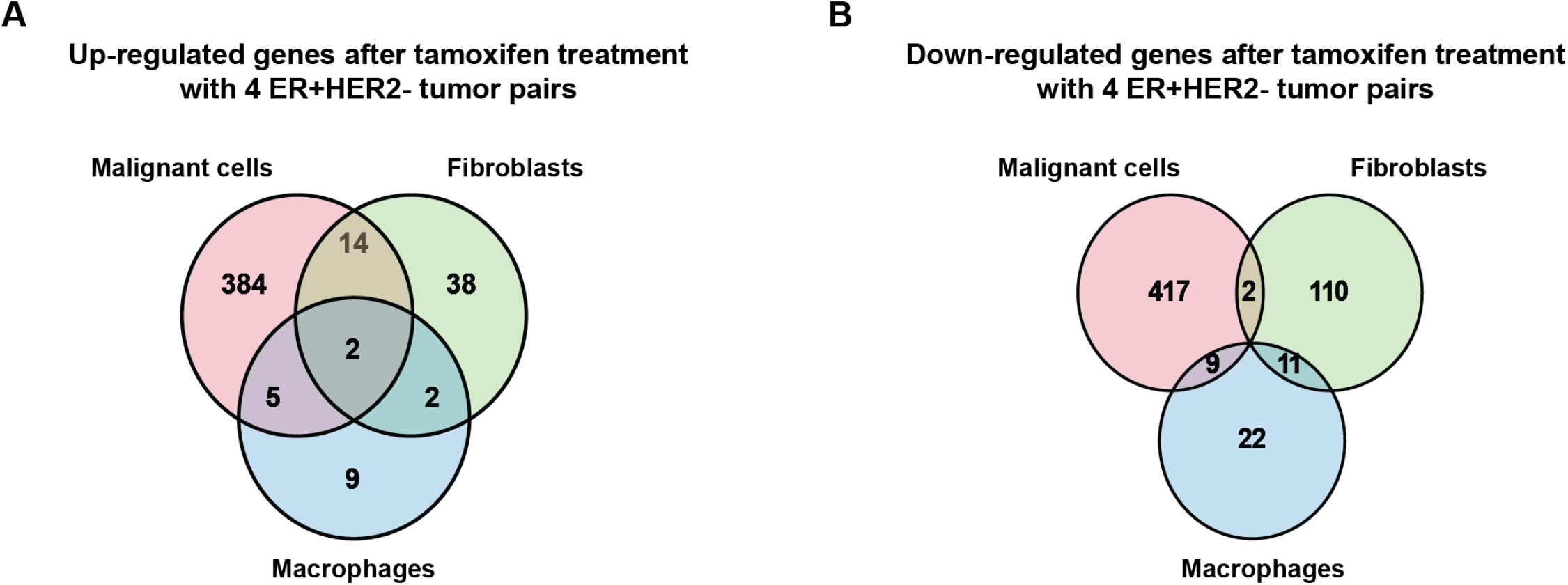
Cell type-specific response to tamoxifen in primary human breast tumors. Venn diagrams of (A) up- and (B) down-regulated genes with tamoxifen treatment by cell type. Most tamoxifen-regulated genes are cell type-specific with minimal overlap across compartments.

**Extended Data Fig. 5.**
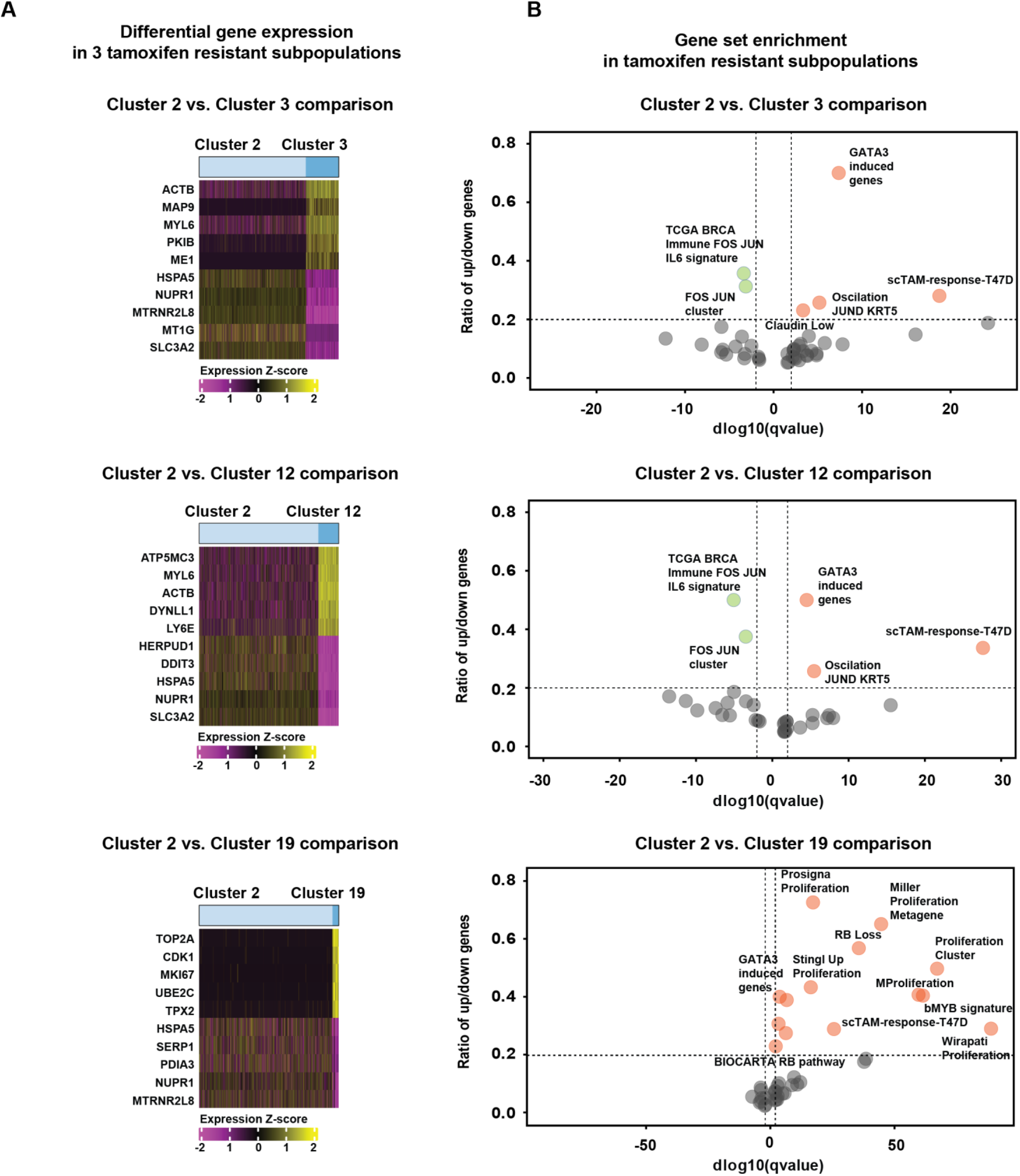
Characteristic gene expression profile of tamoxifen resistant clusters. (A) Heatmaps showing differentially expressed genes between resistant clusters (3,12, 19) and the tamoxifen-sensitive cluster 2. (B) Volcano plot comparing gene set enrichment and depletion of resistant clusters. Significant gene sets were defined as a gene ratio of ≥ 0.2 and a log10qvalue of < −2 or > 2. To simplify visualization, only gene sets with names containing relevant terms were plotted. Supplementary Table 7-9 provides a comprehensive list of the enriched and depleted gene sets for the tamoxifen-resistant clusters (Clusters 3, 12, and 19) compared to the tamoxifen response cluster (Cluster 2).

**Extended Data Fig. 6.**
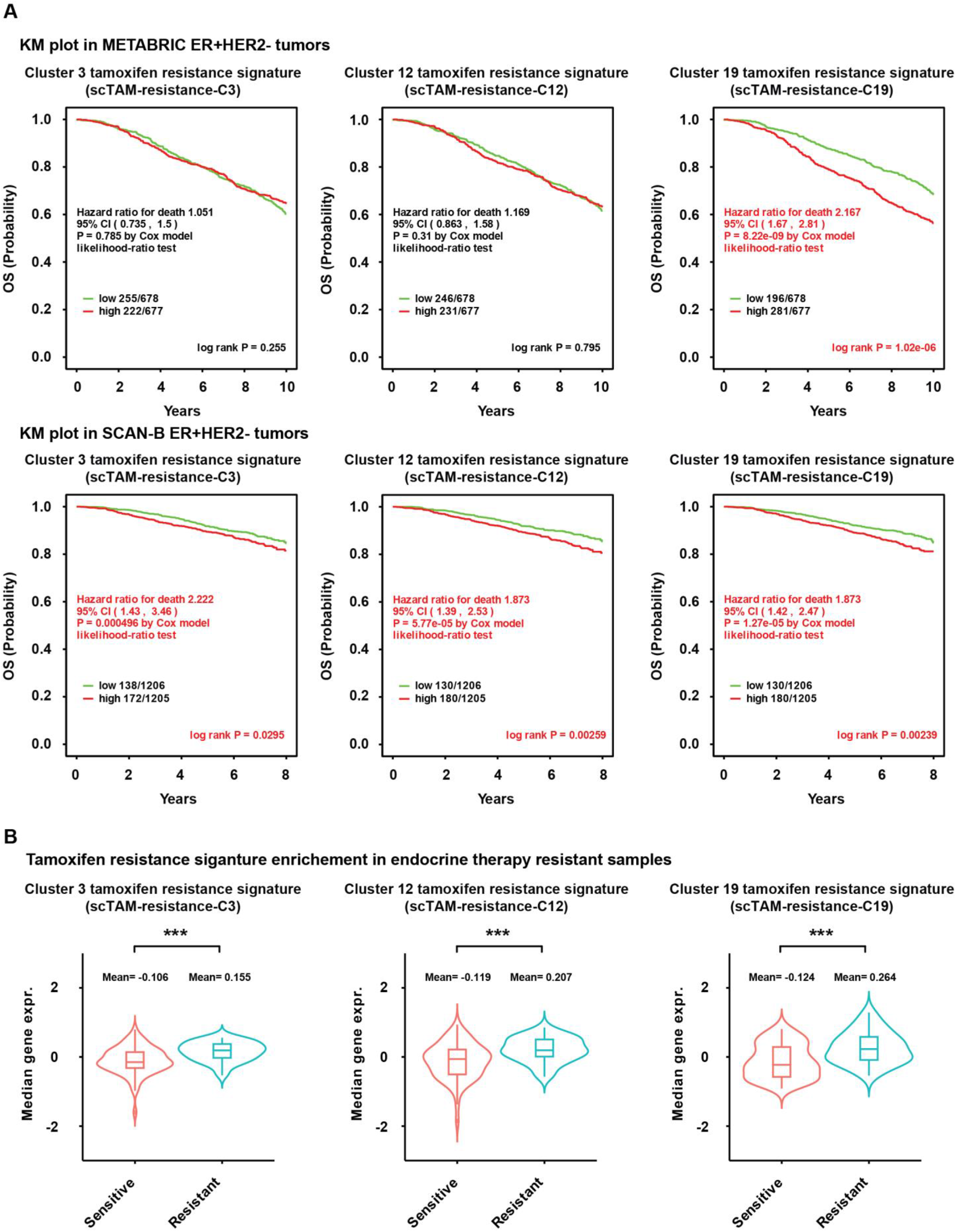
Prognostic signatures of resistant cell populations. (A) Clinically annotated transcriptional data from ER+ patients in the METABRIC and SCAN-B studies was analyzed to evaluate the prognostic significance of resistant cell-specific signatures. Kaplan-Meier (KM) curves demonstrate that signature scores from cluster 3 and cluster 12 were associated with worse survival in SCAN-B but not in METABRIC. The cluster 19 signature was prognostic in METABRIC (HR 2.2, p< 0.001) and SCAN-B (HR1.9, p=0.002). Significance was determined using the log-rank test. (B) We obtained data of clinically annotated endocrine therapy sensitive and resistant tumors from Xia et al^51^. All three tamoxifen resistance signatures were enriched in resistant tumors compared to sensitive. Significance was determined using the Wilcoxon Sum-Rank test, ***p<0.001.

## SUPPLEMENTARY TABLES

**Supplementary Table 1: Comparison of Normal_01 with/without 12hr suspension**

Tab 1. Upregulated genes after 12hr suspension

Tab 2. Enriched Hallmark gene sets for upregulated genes after 12hr suspension

Tab 3. Enriched gene sets from curated breast cancer signatures for upregulated after 12hr suspension

Tab 4. Downregulated genes after 12hr suspension

Tab 5. Enriched Hallmark gene sets after 12hr suspension

Tab 6. Enriched gene sets from curated breast cancer signatures for downregulated genes after 12hr suspension

**Supplementary Table 2: Comparison of tamoxifen and control treated basal epithelial cells (BEp) from normal samples** Tab 1. Upregulated genes with tamoxifen treatment within BEp

Tab 2. Enriched Hallmark gene sets for upregulated genes within BEp

Tab 3. Enriched gene sets from curated breast cancer signatures for upregulated genes within BEp

Tab 4. Downregulated genes with tamoxifen treatment within BEp

Tab 5. Enriched Hallmark gene sets for downregulated genes within BEp

Tab 6. Enriched gene sets from curated breast cancer signatures for downregulated genes within BEp

**Supplementary Table 3: Comparison of tamoxifen-treated and control-treated luminal progenitor cells (LEp_prog) from normal samples**

Tab 1. Upregulated genes with tamoxifen treatment within LEp_prog

Tab 2. Enriched Hallmark gene sets for upregulated genes within LEp_prog

Tab 3. Enriched gene sets from curated breast cancer signatures for upregulated genes within LEp_prog

Tab 4. Downregulated genes with tamoxifen treatment within LEp_prog

Tab 5. Enriched Hallmark gene sets for downregulated genes within LEp_prog

Tab 6. Enriched gene sets from curated breast cancer signatures for downregulated genes within LEp_prog

**Supplementary Table 4: Comparison of tamoxifen and control treated mature luminal cells (LEp) from normal samples** Tab 1. Upregulated genes with tamoxifen treatment within LEp

Tab 2. Enriched Hallmark gene sets for upregulated genes within LEp

Tab 3. Enriched gene sets from curated breast cancer signatures for upregulated genes within LEp

Tab 4. Downregulated genes with tamoxifen treatment within LEp

Tab 5. Enriched Hallmark gene sets for downregulated genes within LEp

Tab 6. Enriched gene sets from curated breast cancer signatures for downregulated genes within LEp

**Supplementary Table 5: T47D response to tamoxifen**

Tab 1. Upregulated genes with tamoxifen treatment within T47D cell line

Tab 2. Enriched Hallmark gene sets for upregulated genes within T47D cell line

Tab 3. Enriched gene sets from curated breast cancer signatures for upregulated genes within T47D cell line

Tab 4. Downregulated genes with tamoxifen treatment within T47D cell line

Tab 5. Enriched Hallmark gene sets for downregulated genes within T47D cell line

Tab 6. Enriched gene sets from curated breast cancer signatures for downregulated genes within T47D cell line

**Supplementary Table 6: Response to tamoxifen in InferCNV+ malignant cells within the selected 4 pairs of ER+/HER2- tumors**

Tab 1. Upregulated genes with tamoxifen treatment

Tab 2. Enriched Hallmark gene sets for upregulated genes

Tab 3. Enriched gene sets from curated breast cancer signatures for upregulated genes

Tab 4. Downregulated genes with tamoxifen treatment

Tab 5. Enriched Hallmark gene sets for downregulated genes

Tab 6. Enriched gene sets from curated breast cancer signatures for downregulated genes

Tab 7. Genes defining scTAM-resistance-M

**Supplementary Table 7 Comparison of tamoxifen and control treated non-malignant cell subpopulations within the selected 4 pairs of ER+/HER2- tumors**

Tab 1. Upregulated genes with tamoxifen treatment within fibroblasts

Tab 2. Enriched Hallmark gene sets for upregulated genes within fibroblasts

Tab 3. Enriched gene sets from curated breast cancer signatures for upregulated genes within fibroblasts

Tab 4. Downregulated genes with tamoxifen treatment within fibroblasts

Tab 5. Enriched Hallmark gene sets for downregulated genes within fibroblasts

Tab 6. Enriched gene sets from curated breast cancer signatures for downregulated genes within fibroblasts

Tab 7. Upregulated genes with tamoxifen treatment within macrophage cells

Tab 8. Enriched Hallmark gene sets for upregulated genes within macrophage cells

Tab 9. Enriched gene sets from curated breast cancer signatures for upregulated genes within macrophage cells

Tab 10. Downregulated genes with tamoxifen treatment within macrophage cells

Tab 11. Enriched Hallmark gene sets for downregulated genes within macrophage cells

Tab 12. Enriched gene sets from curated breast cancer signatures for downregulated genes within macrophage cells

**Supplementary Table 8: Comparison of upregulated and downregulated genes in resistant cell subpopulation cluster 3 compared to cluster 2**

Tab 1. Upregulated genes with tamoxifen treatment within cluster 3 compared to cluster 2

Tab 2. Enriched Hallmark gene sets for upregulated genes within cluster 3 compared to cluster 2

Tab 3. Enriched gene sets from curated breast cancer signatures for upregulated genes within cluster 3 compared to cluster 2

Tab 4. Downregulated genes with tamoxifen treatment within cluster 3 compared to cluster 2

Tab 5. Enriched Hallmark gene sets for downregulated genes within cluster 3 compared to cluster 2

Tab 6. Enriched gene sets from curated breast cancer signatures for downregulated genes within cluster 3 compared to cluster 2

Tab 7. Genes defining scTAM-resistance-C3

**Supplementary Table 9: Comparison of upregulated and downregulated genes in resistant cell subpopulation cluster 12 compared to cluster 2**

Tab 1. Upregulated genes with tamoxifen treatment within cluster 12 compared to cluster 2

Tab 2. Enriched Hallmark gene sets for upregulated genes within cluster 12 compared to cluster 2

Tab 3. Enriched gene sets from curated breast cancer signatures for upregulated genes within cluster 12 compared to cluster 2

Tab 4. Downregulated genes with tamoxifen treatment within cluster 12 compared to cluster 2

Tab 5. Enriched Hallmark gene sets for downregulated genes within cluster 12 compared to cluster 2

Tab 6. Enriched gene sets from curated breast cancer signatures for downregulated genes within cluster 12 compared to cluster 2

Tab 7. Genes defining scTAM-resistance-C12

**Supplementary Table 10: Comparison of upregulated and downregulated genes in resistant cell subpopulation cluster 19 compared to cluster 2**

Tab 1. Upregulated genes with tamoxifen treatment within cluster 19 compared to cluster 2

Tab 2. Enriched Hallmark gene sets for upregulated genes within cluster 19 compared to cluster 2

Tab 3. Enriched gene sets from curated breast cancer signatures for upregulated genes within cluster 19 compared to cluster 2

Tab 4. Downregulated genes with tamoxifen treatment within cluster 19 compared to cluster 2

Tab 5. Enriched Hallmark gene sets for downregulated genes within cluster 19 compared to cluster 2

Tab 6. Enriched gene sets from curated breast cancer signatures for downregulated genes within cluster 19 compared to cluster 2

Tab 7. Genes defining scTAM-resistance-C19

